# Resting-state aperiodic neural dynamics predict individual differences in visuomotor performance and learning

**DOI:** 10.1101/2021.04.08.438941

**Authors:** Maarten A. Immink, Zachariah R. Cross, Alex Chatburn, James Baumeister, Matthias Schlesewsky, Ina Bornkessel-Schlesewsky

**Affiliations:** Sport, Health, Activity, Performance and Exercise (SHAPE) Research Centre, Flinders University, Adelaide, Australia; Cognitive and Systems Neuroscience Research Hub, University of South Australia, Adelaide, Australia; Australian Research Centre for Interactive and Virtual Environments, University of South Australia, Adelaide, Australia

**Keywords:** visuomotor performance, reaction time, perception, individual differences, motor learning, EEG, aperiodic activity, 1/f brain activity

## Abstract

An emerging body of work has demonstrated that resting-state non-oscillatory, or aperiodic, 1/*f* neural activity is a functional and behaviorally relevant marker of cognitive function capacity. In the motor domain, previous work has only applied 1/*f* analyses to description of action coordination and performance. The value of aperiodic resting-state neural dynamics as a marker of individual visuomotor performance capacity remains unknown. Accordingly, the aim of this work was to investigate if individual 1/*f* intercept and slope parameters of aperiodic resting-state neural activity predict reaction time and perceptual sensitivity in an immersive virtual reality marksmanship task. The marksmanship task required speeded selection of target stimuli and avoidance of non-target stimuli selection. Motor and perceptual demands were incrementally increased across task blocks and participants performed the task across three training sessions spanning one week. When motor demands were high, steeper individual 1/*f* slope predicted shorter reaction time. This relationship did not change with practice. Increased 1/*f* intercept and a steeper 1/*f* slope were associated with higher perceptual sensitivity, measured as *d’*. However, this association was only observed under the highest levels of perceptual demand and only in the initial exposure to these conditions. Individuals with a lower 1/*f* intercept and a shallower 1/*f* slope demonstrated the greatest gains in perceptual sensitivity from task practice. These findings demonstrate that individual differences in motor and perceptual performance can be accounted for with resting-state aperiodic neural dynamics. The 1/*f* aperiodic parameters are most informative in predicting visuomotor performance under complex and demanding task conditions. In addition to predicting capacity for high visuomotor performance with a novel task, 1/*f* aperiodic parameters might also be useful in predicting which individuals might derive the most improvements from practice.

## Introduction

The identification of individual characteristics that predict motor skill learning and performance proficiency continues to be a central practical and theoretical issue. On the practical side, individual predictors allow the practitioner to identify individuals with high capacity for learning and performance of a motor skill. Predictors are also valuable to inform individualization of training environments to enhance future performance outcomes. Theoretically, the identification of individual predictors affords the advancement of our understanding of the essential requirements to acquire and perform a class of actions. Moreover, as action is inherently noisy due to inter-individual variance, the identification of individual predictors offers utility in accounting for this noise such that the signal – the learning or performance outcome -can be more directly considered.

Historically, under the rubric of motor abilities, the search for individual predictors of skilled performance initially concentrated on sensorimotor characteristics (e.g., Adams, 1957; Fleishman, 1960; Welch & Henry, 1971; Fleishman, 1972; Fleischman & Mumford, 1989). Subsequent work (e.g., Ackerman, 1987, 1988; Ackerman & Cianciolo, 2000) has since highlighted differences in cognitive function as key individual determinants of motor skill acquisition and performance. For example, working memory capacity (Christou et al., 2016) and metacognitive ability (Sinanaj et al., 2015) have both been shown to predict learning and performance of visuomotor skills. Since these characteristics generally predict proficiency in a range of tasks including those that assess fluid intelligence (Kyllonen & Christal, 1990; Swanson & McMurran, 2018), their association with visuomotor performance could more globally reflect how action arises from a complex self-organizing system (Van Orden et al., 2003). Executive function and action performance arise from the activation of a number of shared brain regions including the prefrontal cortex, cingulate cortex, premotor cortices, motor cortex, parietal cortex and the basal ganglia (Mirabella, 2014; Immink et al., 2020). This overlap suggests that higher order cognitive function such as working memory and metacognition might predict visuomotor performance because they share self-organizing properties of the brain. As such, it would seem logical to identify self-organizing properties of the brain that can be utilized as biological markers for individual capacity in the learning and performance of visuomotor skills.

There have been several previous attempts to quantify neurophysiological markers of motor learning and performance (Baetu et al., 2015; Andreska et al., 2020; Herszage et al., 2020). Such attempts have included resting-state electroencephalography (EEG) activity, which has been shown to predict visuomotor performance with the rotary pursuit task (Wu et al., 2014). These findings suggest that resting-state EEG profiles might offer promising insights into assessing individual self-organizing neural characteristics as predictors of visuomotor capacity (Cheron et al., 2016). This particularly so given the accessibility of undertaking resting EEG recording both in laboratory and practical settings as well other advantages of EEG over other neural recording methodology including temporal resolution and simultaneous recording of multiple cortical sites. However, one key consideration is the identification of the resting-state EEG measure that most reliably predicts skill performance and learning.

Electrical neural activity is comprised of broadband scale-free (otherwise referred to as fractal or aperiodic) and oscillatory activity (He et al., 2010; He, 2014). Research has traditionally ignored scale-free activity, treating it as nuisance “noise” (Groppe et al., 2013), and instead focused on extracting oscillations (i.e., in the delta, theta, alpha and beta band frequencies) from resting-state and task-related recordings, such as during perceptual decision-making tasks. While this research has revealed that activity in certain frequency ranges (e.g., the alpha band; 8 – 12 Hz) is correlated with a range of behavioral, perceptual and cognitive outcomes (Cheron et al., 2016), recent work has demonstrated that aperiodic activity better predicts processing speed than resting alpha oscillatory activity (Ouyang et al., 2020).

The aperiodic component of electrophysiological signals is distinguishable from oscillatory activity, manifesting as a straight line on a power spectral density (PSD) plot (Donoghue et al., 2020; see Figure 1 for a visualization of aperiodic and oscillatory components of a PSD plot), which follows the 1/*f* power-law exponent (He, 2014). The 1/*f* power-law exponent refers to the observation that power is highest at lower frequencies, with power exponentially decreasing with increasing frequency (Demuru & Fraschini, 2020; Donoghue et al., 2020). The power-law exponent, or β, reflects the steepness of the 1/*f* slope, with higher β values indicating a steeper slope and lower values reflecting a shallower (i.e., flatter) slope (He, 2014). Mechanistically, the 1/*f* slope is argued to reflect an excitation-inhibition balance in recurrent neural networks which maximises information processing capacities (Lendner et al., 2020; Weber et al., 2020), with steeper slopes indexing enhanced neural inhibition and vice versa. Aperiodic activity can also be quantified by its spectral offset (i.e., the intercept), with a higher intercept hypothesized to be reflective of increased neural population spiking (Manning et al., 2009; Miller et al., 2012), and is positively correlated with the blood-oxygen-level-dependent (BOLD) signal (Winawer et al., 2013; Jacob et al., 2021) obtained from functional magnetic resonance imaging (fMRI).

**Fig. 1.**
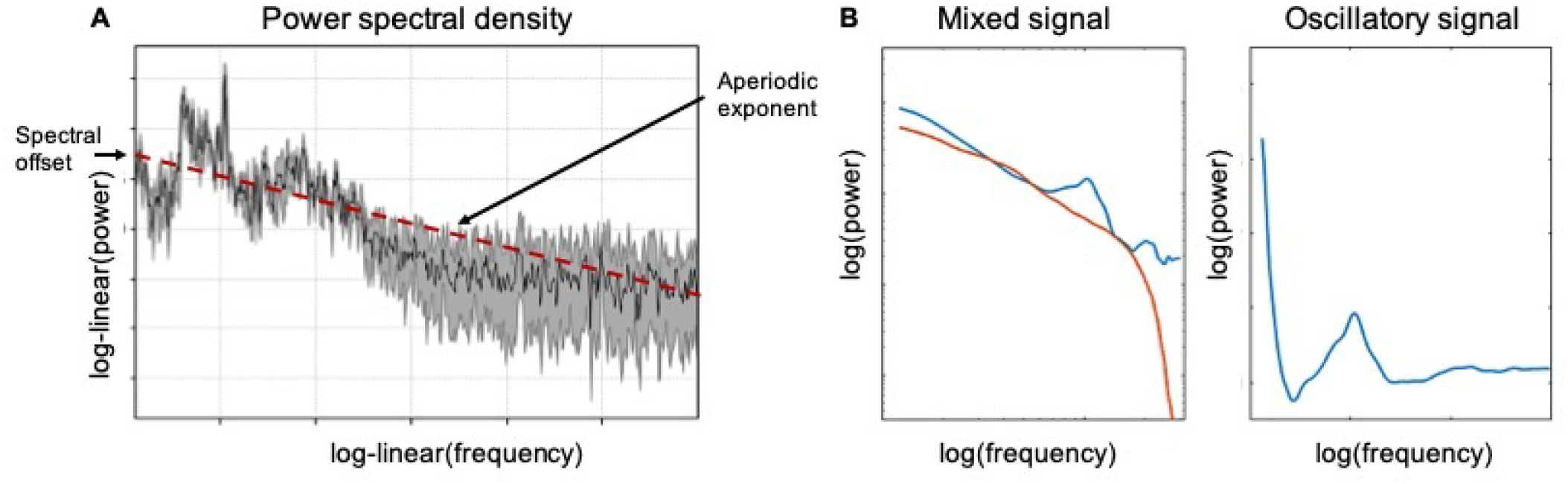
Power spectral density (PSD) estimates of fractal (aperiodic 1/*f*) and oscillatory activity from a single individual at channel Pz during a memory task. (A) PSD plot illustrating the spectral offset (y-axis) and aperiodic exponent (dashed red line). (B) The PSD plot on the right reflects the difference wave between the total (blue) and fractal (red) PSD estimates that are illustrated in the PSD plot on the left. Log transformed power (i.e., strength of neural activity) is represented on the y-axis (higher values indicate greater activity), while frequency (Hz; cycles per second) is represented on the x-axis (values further to the right indicate higher frequency).

**Fig. 2.**
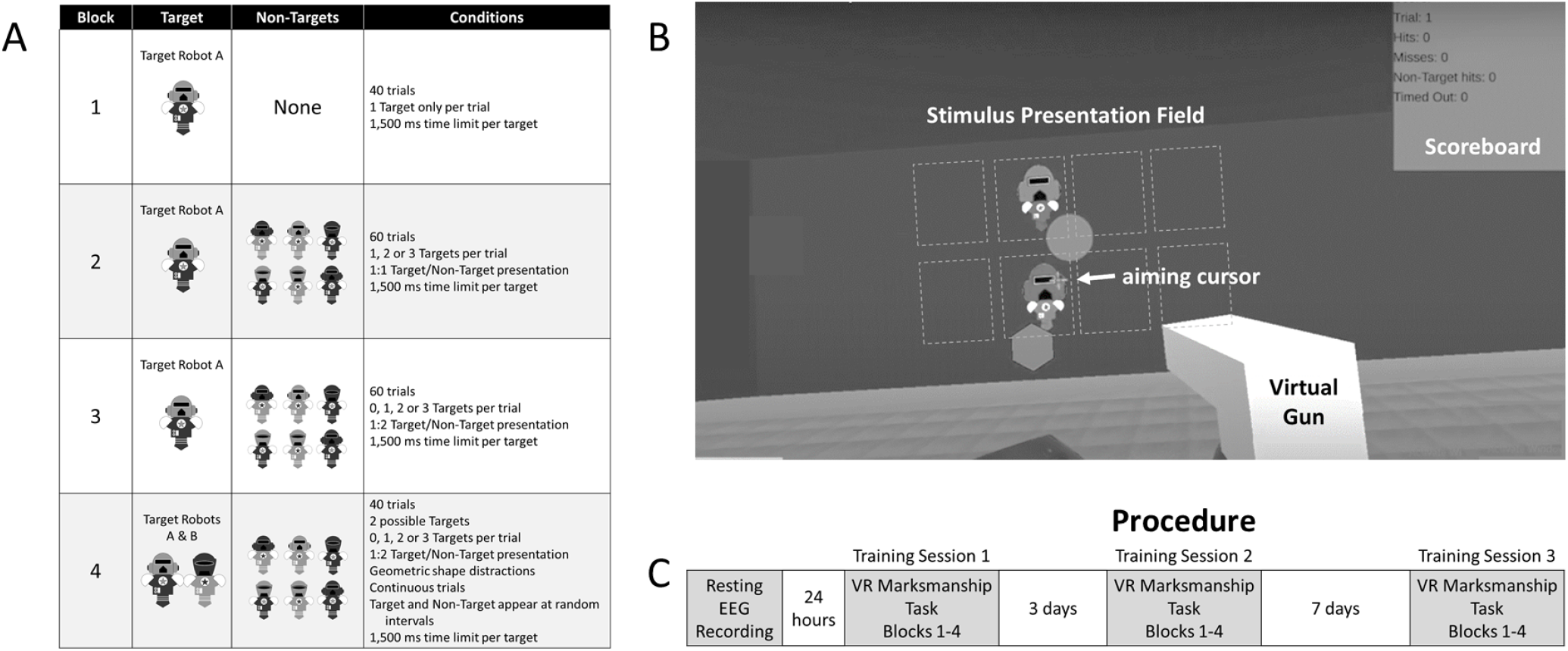
(A) Conditions for Blocks 1 to 4 of the Virtual Reality (VR) Marksmanship Task. The target and non-target robots were equivalent in size in the VR environment. (B) Immersive VR environment for the gameified Marksmanship Task. The virtual gun was controlled by a hand controller in the right hand allowing for aiming with a displayed cursor and selection of stimuli. The environment view changed with head movement. Note: The squares were not presented in the environment. They are included to illustrate the 8 possible locations that stimuli were presented. (C) Procedure for recording resting EEG followed by three training sessions with the VR Marksmanship Task.

Estimation of aperiodic activity from resting-state EEG recordings is straightforward (e.g., Demuru et al., 2020; Donoghue et al., 2020; Ouyang et al., 2020) resulting in an accessible measure of resting-state derived neural activity that can index intrinsic network-related activity with high test re-test reliability. The 1/*f* aperiodic parameters (i.e., spectral slope and intercept) appear to be robust markers of neural information processing across a range of domains. Indeed, an emerging body of work has revealed the effectiveness of 1/*f* in predicting individual capacity for processing speed (Ouyang et al., 2020) and artificial grammar learning (Cross et al., 2020). Specific to the motor domain, there have been earlier applications of 1/*f* (also referred to as pink noise) scaling to describe self-organizing properties of skilled movement performance (Diniz et al., 2011) including reaction time (Gilden et al., 1995; Clayton & Frey, 1997), movement timing (Wijnants et al., 2009), and oscillation of finger tapping (Rigoli et al., 2014) and music instrument performance (Colley & Dean, 2019). However, there has been no previous work that has addressed 1/*f* aperiodic resting-state EEG activity as a predictor of visuomotor performance.

The aim of the present work was to investigate if the 1/*f* intercept and slope estimated from resting-state EEG predict motor and perceptual dimensions of visuomotor performance. Given that the 1/*f* intercept reflects mean neural population spiking and, by extension, higher broadband power (Donoghue et al., 2020), a higher resting-state intercept would be expected to predict both motor and perceptual performance. Further, as a steeper 1/*f* slope indexes greater neural excitation-inhibition balance to the extent of maximising information processing (Lendner et al., 2020; Weber et al., 2020), then a steeper resting-state slope would predict greater capacity for visuomotor processing speed and perceptual sensitivity to target information. We also investigated if individual 1/*f* intercept and slope predicted performance over increased task difficulty by including four task blocks were target selection was rendered more complex by the presence of multiple targets as well as non-targets. Finally, we investigated how task training influenced performance prediction from individual 1/*f* intercept and slope over a 1-week period of training involving three training sessions.

## Methods

### Participants

Forty-five adults (Mage = 22.67 ± 3.85 years, 23 females) volunteered to participated in the Experiment. Participant eligibility included: 18-35 years of age; right-hand dominance based on the Edinburgh Handedness Inventory -Short Form (Veale, 2014); normal or corrected vision; no sensory, motor or cognitive impairments; no history of psychiatric disorders. Participants were recruited from the Adelaide metropolitan community in Australia and were naïve to the aim of the Experiment. All participants provided written informed consent prior to participation. The Experiment protocol was approved by the University of South Australia Human Research Ethics Committee.

### Tasks and apparatus

#### Virtual reality visual-motor marksmanship task

A customized virtual reality (VR) visual-motor marksmanship task was developed and implemented within the Unity game engine platform (https://unity.com/). A head-mounted HTC VIVE Pro (HTC Inc, Bellevue, Washington) displayed the VR and a VIVE Controller held in the right hand was used to aim and shoot, via trigger pull, visual targets in the VR environment. Movement of the headset and hand-held controller was tracked with two VIVE base stations. Participants completed the task while seated.

The marksmanship task involved the presentation of cartoon robots that were distinguished by their helmet shape as well as red or blue coloring of the helmet and armor. Robots appeared in a two by four matrix of spatial regions that surrounded a central point in the VR environment. The matrix was comprised of a row above and a row below the central point and two columns to the left and right of the central point. Participants aimed and shot at the central point to commence a trial, after which robots were presented in the spatial locations. Aiming was assisted by the presentation of an aiming cursor (cross-hairs) in the VR environment. Gamification of the marksmanship task involved a scoring system where shooting a target robot was awarded points, but this award depreciated with target hit latency up to 1,500 milliseconds. Score feedback for shooting a robot was briefly displayed and a running score board was displayed in the left-hand side of the VR environment.

The marksmanship task involved performance of four blocks representing increasing difficulty in terms of the number of target and non-target robots presented in a trial. Block 1 was designed to assess proficiency in detecting, aiming and selecting a visual target by presenting only one target robot, referred to as target robot A, in each of the 40 trials. Participants were instructed to shoot the target as fast possible. If the target was not shot within the 1,500 ms time limit, as miss result was recorded and a “time out” message was displayed. Following Block 1, the subsequent blocks were designed to introduce and increase demands on selective attention, perceptual discrimination, and response inhibition through the introduction of non-target robots. In Blocks 2 to 4, if a non-target robot was shot, an error message was presented, penalty points were applied and a false alarm response was recorded. In each of the 60 trials under Block 2, one, two or three target robot A were presented along with non-target robots at a 1:1 ratio. Targets and non-targets were presented at once in a trial. Three variations of non-target robots differed from target robot A in terms of either the helmet shape or helmet and armor color. In Block 3, involving 60 trials, presented target robot A and non-target robots at a ratio of 1:2. In addition, Block 3 included catch-trials involving the presentation of only non-target robots. Targets and non-targets were presented at once in each trial of Block 3. For Block 4, involving 40 trials, a new target robot, termed target robot B, was presented in addition to target robot A and the non-target robots. The new target robot differed from target robot A with respect to both the helmet shape, and helmet and armor color. In Block 4, trials were conducted continuously such that target and non-target robots were presented at random intervals throughout the block with multiple robots present in the VR environment simultaneously. Further visual distraction was introduced in Block 4 with colored static and moving geometric shapes appearing in the continuous trials.

The motor component of marksmanship performance in Blocks 1 to 4 of the task was based on reaction time (RT). RT was measured in milliseconds as the latency between target robot presentation and selection of the target (e.g., shooting the target with the hand controller). Mean RT was calculated for each block based on all instances that a target robot was selected within the 1,500 ms time limit. The perceptual component of task performance in each block was based on Hit rate, the proportion of presented targets that were selected, and False Alarm (FA) rate, the proportion of non-targets that were selected. For Blocks 2 to 4, perceptual sensitivity was based on d prime (*d’*), which was calculated for each participant from the Hit and FA rate in each block (Swets et al., 1961). Hit and FA rates were transformed into z-normalized ZHit and ZFA distributions and then d’ for the block was calculated (Macmillan & Creelman, 1990) based on the formula:

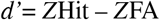

*d’* represents the distance between the probability distributions of Hits and FA. Higher perceptual sensitivity is reflected by higher *d’* values reflecting maximization of hits (selecting target robots) and false alarm minimization (selecting non-target robots).

### EEG Recording and Preprocessing

EEG data were recorded at rest using a LiveAmp (Brain Products GmbH, Gilching, Germany) with 32 active Ag/AgCl electrodes mounted in an elastic cap (Brain Cap, Brain Products GmbH, Gilching, Germany). Electrode placement followed the 10/20 system. The EEG was recorded with a sampling rate of 500 Hz, referenced to FCz with the ground electrode positioned at AFz. Impedance for each electrode was kept below 5 kOhm during the recording. Electrooculogram (EOG) recordings were obtained from below the left eye and from the outer canthus of the right eye.

The EEG data were analyzed using MNE-Python (Gramfort et al., 2013). EEG data were re-referenced offline to the average of TP9 and TP10 and filtered with a digital phase-true finite response (FIR) band-pass filter from 0.1 -40 Hz to remove slow signal drifts and high frequency activity.

#### Quantification of aperiodic neural dynamics

In order to estimate the 1/*f* power-law exponent characteristic of background spectral activity, we used the irregular-resampling auto-spectral analysis method (IRASA; Wen & Liu, 2015), implemented in the YASA toolbox in MNEPython (Vallat, 2019). The IRASA method (Wen et al., 2015; Wen & Liu, 2016) separates the fractal and oscillatory components in the power spectrum of the EEG. Briefly, IRASA computes the original power spectral density (PSD) and resamples the EEG by multiple non-integer factors and their reciprocals (i.e., h and their reciprocals 1/h). For each set of signals that have been resampled, the PSD is calculated and the geometric mean of both is taken, with the power associated with the PSD being redistributed away from its original frequencies by a frequency offset that varies by the resampled factor. By contrast, the power associated with the fractal (i.e., 1/*f*) component remains the same power-law statistical distribution irrespective of the resampling factor. By taking the median of the resampled PSD estimates, the power spectrum of the fractal component can then be taken, with the difference between the original PSD and the extracted fractal estimate serving as a proxy for estimated power of the oscillatory component. The aperiodic signal, L, is modeled using an exponential function in semilog-power space (linear frequencies and log PSD) as:

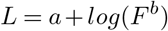

where a is the intercept, b is the slope, and F the vector of input frequencies. For more information on the IRASA method, please refer to Wen et al. (2016). For a schematic illustration of oscillatory versus aperiodic components estimated from resting-state EEG, see Figure 3.

### Procedure

Participants first attended a laboratory to record resting EEG for the purpose of measuring resting-state 1/*f* dynamics. Participants were fitted with the EEG cap, prepared for EEG recording and then completed a 5-minute resting eyes open, eyes closed protocol. About 24 hours later, participants returned for their first training session with the VR marksmanship task. Participants were fitted with the VR headset, provided the hand controller, and then completed Blocks 1 to 4 of the task. Training session 2 was completed approximately 72 hours after training session 1 and training session 3 was completed 7 days after training session 2. Training sessions 2 and 3 were conducted similarly to training session 1. No task training was completed between sessions.

### Data Analysis

Data were imported into R version 4.0.2 (R Core Team, 2020) and analysed using linear mixed effects models fit by restricted maximum likelihood (REML) estimates using lme4 (Bates et al., 2015). Models examining RT as the outcome variable took the following form:

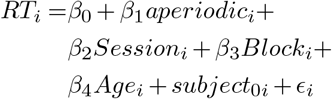

Models examining sensitivity (i.e., *d’*) to target stimuli included the same predictor variables, taking the following form:

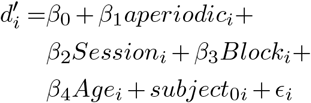

where, in both models, aperiodic refers to either the 1/*f* slope or intercept, session encodes sessions 1 – 3, block refers to blocks 1 – 4. Random effects on the intercept for Subject was included to account for systematic variance amongst participants (Van Dongen et al., 2004; Judd et al., 2012), while age was added as a main effect to control for any age-related effects of aperiodic activity on behavior (e.g., Voytek et al., 2015). Asterisks denote interaction terms; pluses denote additive terms. Type II Wald χ2-tests from the car package (Fox & Weisberg, 2019) were used to provide p-value estimates, while effects were plotted using the package effects (Fox et al., 2019) and ggplot2 (Wickham, 2016). Session and Block were specified as ordered factors and were examined using polynomial contrasting. We also used an 83% confidence interval (CI) threshold, as this approach corresponds to the 5% significance level with non-overlapping estimates (Austin & Hux, 2002).

## Results

### Task difficulty and training session effects

Mixed effects modelling of RT revealed significant main effects of Block (χ2(3) = 6596.10, *p* < .001), Session (χ2(2) = 221.55, *p* < .001), and a significant Block x Session interaction (χ2(6) = 17.34, *p* = .008), see Table 1. As illustrated in Figure 4A, across training sessions, the shortest RT was in Block 1. While RT did not significantly differ between Block 3 and 4 across sessions, Block 2 RT was significantly shorter than Block 3 and 4 RT in Session 1 and 3.

**Table 1.** Target Reaction Time, Hit Rate, False Alarm Rate and *d’* sensitivity index for Block conditions in the virtual reality marksmanship task. Hit Rate was the ratio of visual targets selected relative to those presented in a block. The False Alarm Rate, the ratio of non-targets selected in proportionally to those presented, is only relevant to Blocks 2 to 4 as in Block 1, only targets were presented. Note. Parentheses reflect the standard error of the mean.

|  | Block | Session |  |  |
| --- | --- | --- | --- | --- |
|  |  | 1 | 2 | 3 |
| Target Reaction Time (ms) | 1 | 560.92 (8.93) | 539.20 (7.55) | 516.09 (6.82) |
|  | 2 | 964.15 (14.24) | 906.48 (12.71) | 869.18 (11.14) |
|  | 3 | 1025.04 (13.09) | 974.87 (13.77) | 941.19 (11.65) |
|  | 4 | 1010.93 (10.88) | 943.66 (11.28) | 915.50 (10.06) |
| Hit Rate | 1 | 0.994 (0.002) | 0.998 (0.001) | 0.999 (0.001) |
|  | 2 | 0.974 (0.003) | 0.986 (0.001) | 0.989 (0.000) |
|  | 3 | 0.924 (0.007) | 0.954 (0.007) | 0.967 (0.005) |
|  | 4 | 0.825 (0.021) | 0.900 (0.016) | 0.908 (0.013) |
| False Alarm Rate | 2 | 0.012 (0.0013) | 0.009 (0.000) | 0.021 (0.005) |
|  | 3 | 0.015 (0.0018) | 0.010 (0.001) | 0.010 (0.001) |
|  | 4 | 0.134 (0.017) | 0.067 (0.013) | 0.054 (0.011) |
| Sensitivity Index ($d'$) | 2 | 4.37 (0.06) | 4.62 (0.03) | 4.53 (0.06) |
|  | 3 | 3.81 (0.09) | 4.24 (0.09) | 4.40 (0.08) |
|  | 4 | 2.45 (0.17) | 3.28 (0.14) | 3.42 (0.14) |

**Fig. 4.**
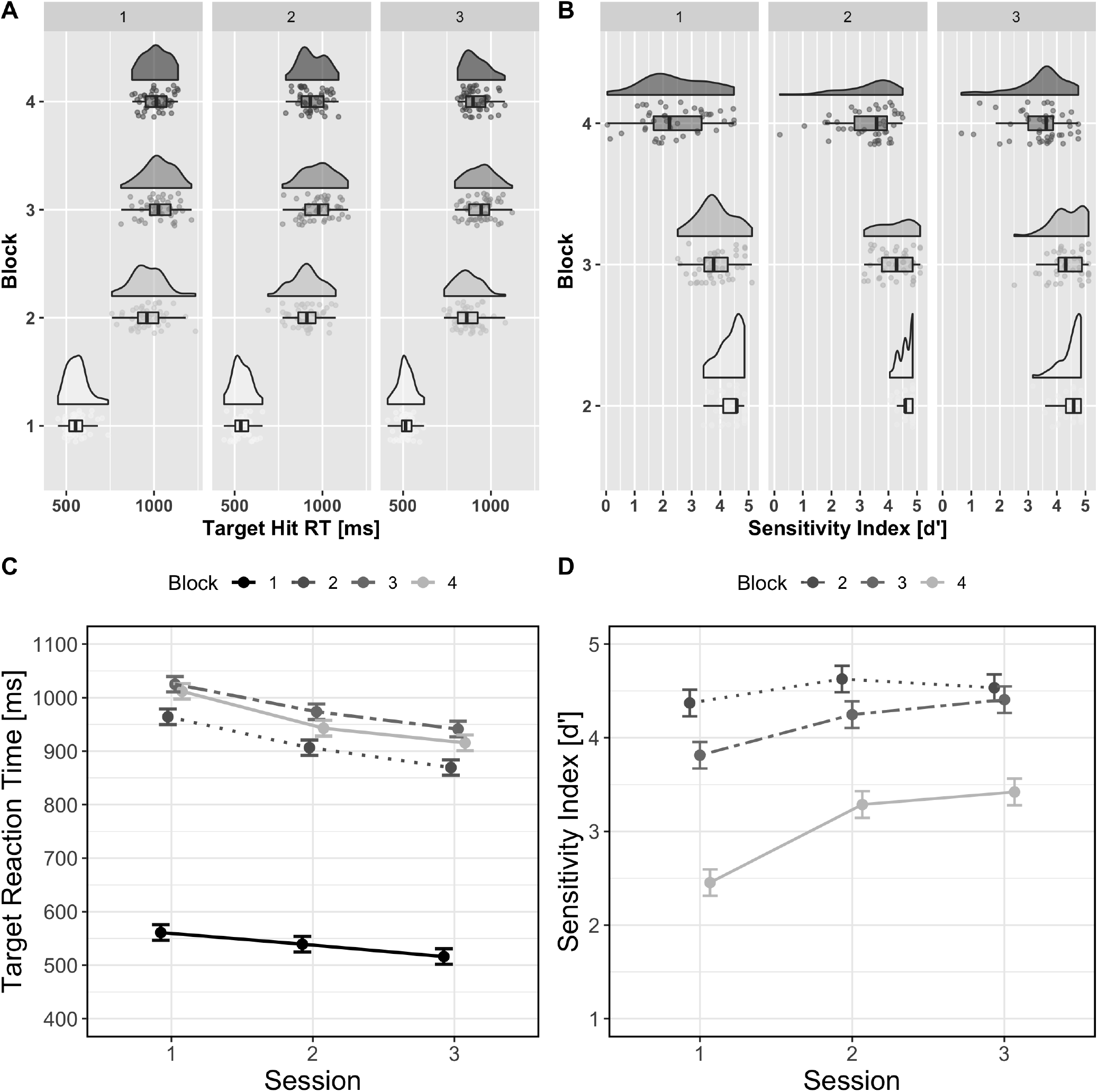
Visualization of the distribution in RT and *d’* across Session and Block. (A) Mean reaction time (ms; y-axis) for target stimuli across Block 1 – 4 (y-axis) and Session (session 1 = left, session 2 = middle, session 3 = right). (B) Mean *d’* scores (x-axis) across Blocks 2 to 3 (y-axis) and Sessions. Data points represent individual participants .(C) Modelled effects of RT (y-axis) across Session (x-axis) for Blocks 1 – 4. Modelled effects of *d’* (y-axis) across Session (x-axis) for Blocks 1 – 4. error bars represent the 83% confidence interval.

**Fig. 3.**
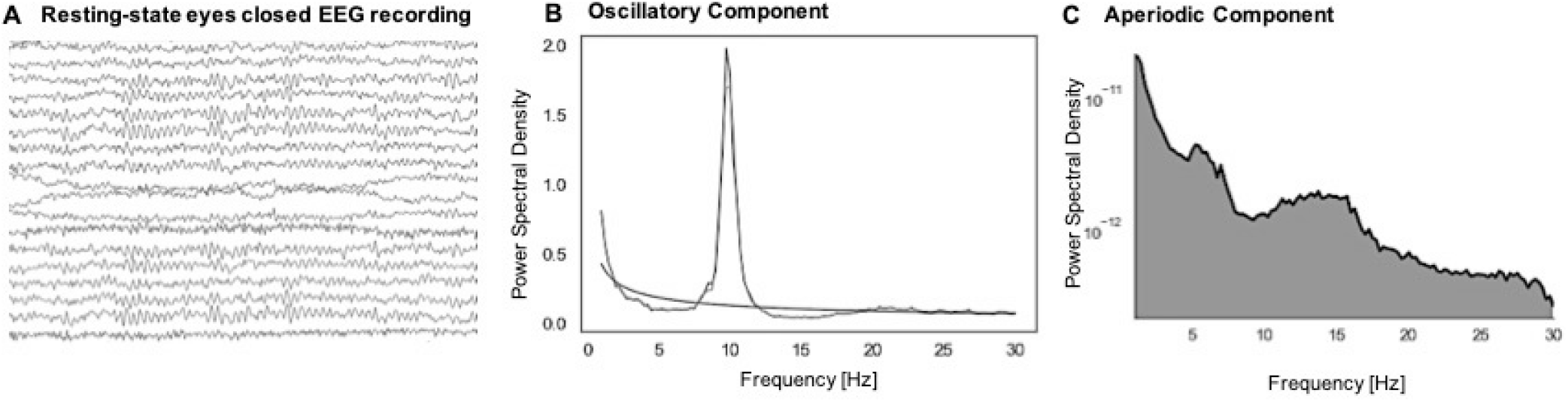
(A) Schematic illustration of a raw EEG trace during a resting-state eyes closed period. (B) Oscillatory component extracted from the EEG. Note the clear spectral peak at roughly 10 Hz, which corresponds to this participant’s individual alpha frequency. (C) Fractal (i.e., aperiodic, 1/*f*) component extracted from the EEG. Note how power (y-axis) decreases exponentially as frequency (x-axis) increases.

Modelling of *d’* revealed significant main effects of Block (χ2(2) = 410.81, *p* < .001) and Session (χ2(2) = 70.40, *p* < .001), as well as a significant Block x Session interaction (χ2(4) = 21.33, *p* < .001), see Table 1. As shown in Figure 4B, d’ scores were highest in Block 2 and then decreased through Blocks 3 and 4. In addition, Block 3 d’ increased from Session 2 to 3, reaching equivalent d’ levels to Block 2. Block 4 d’ also increased across Sessions 1 to 3 but remained lower than Blocks 2 and 4.

### Resting EEG aperiodic neural dynamics as individual predictors of performance

We now report on results from mixed effects modelling of RT and *d’* based on inclusion of EEG aperiodic dynamics as individual predictors of RT and d’ performance across Blocks and Sessions. Overall, the sample mean for 1/*f* intercept was -23.32 (SD = 0.55; range = -24.55 to -21.81) while mean 1/*f* slope was -2.06 (SD = 0.13; range = -2.48 to -1.82). For RT, modelling did not reveal a significant main effect of the 1/*f* intercept (χ2(1) = 1.57, *p* = .20), nor any other significant higher-order interactions with the 1/*f* intercept (see Table S1 in the supplementary material for full model summaries). In contrast, the model examining individual 1/*f* slope indicated a significant 1/*f* slope x Block interaction (χ2(3) = 9.63, *p* = .02). As is illustrated in Figure 5D, a steeper 1/*f* slope (indicated by more negative values on the x-axis) predicted shorter RT in Blocks 2 and 3 but did not predict RT in Blocks 1 and 4 (for full model summaries, see Supplementary Material). For *d’*, there was a significant 1/*f* intercept x Block x Session interaction (χ2(4) = 13.18, *p* = .01). As is illustrated in Figure 5E, a higher 1/*f* intercept was associated with higher *d’* scores in Block 4 within Session 1, but the advantage of higher 1/*f* intercept reduced for Block 4 *d’* in Sessions 2 and In contrast, those with lower 1/*f* intercept demonstrated increased *d’* in Block 4 in Sessions 2 and 3. Examination of individual 1/*f* slope prediction of *d’* indicated a significant 1/*f* slope x Session interaction (χ2(2) = 6.12, *p* = .04). As illustrated in Figure 5F, in Session 1, a steeper 1/*f* slope predicted higher *d’* across task blocks in Session 1. However, in Sessions 2 and 3, the pattern appeared to reverse such that shallower 1/*f* slope was associated with higher *d’*.

**Fig. 5.**
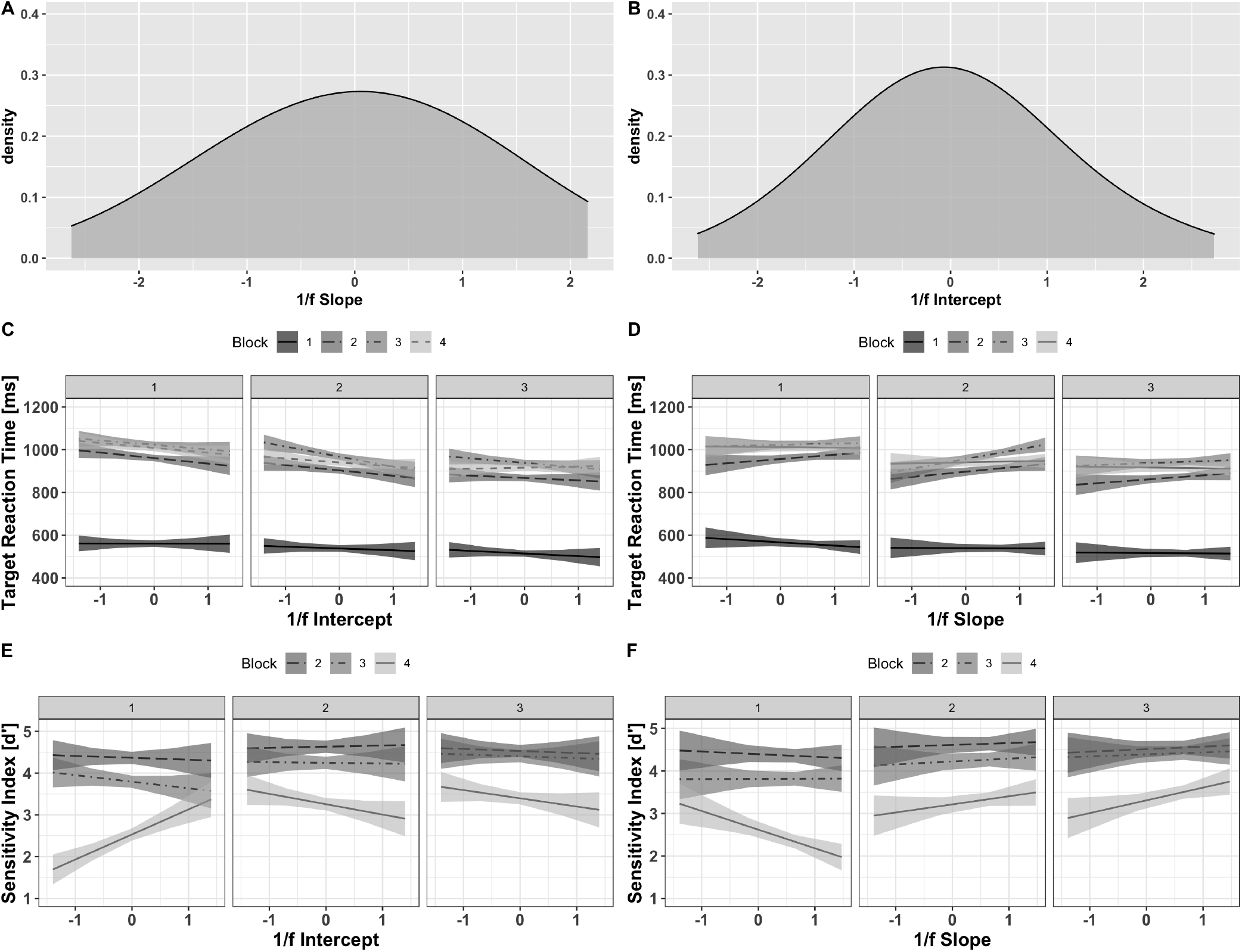
Density plots illustrating the distribution of scaled 1/*f* slope (A) and intercept (B) estimates. (C) Relationship between the 1/*f* intercept (x-axis; less negative values indicate a higher intercept) and RT (y-axis; higher values indicate slower RT) across Session (1 = session 1, 2 = session 2, 3 = session 3) and Block (block 1 = dark solid line, block 2 = lighter dashed line, block 3 = dashed gray line, block 4 = dashed light gray line). (B) Relationship between the 1/*f* slope (x-axis; more negative values indicate a steeper slope) and RT. (C) Relationship between the 1/*f* intercept (x-axis) and *d’* (y-axis; higher values indicate greater sensitivity) across Session (1 = session 1, 2 = session 2, 3 = session 3) and Block. (D) Relationship between the 1/*f* slope (x-axis) and *d’* across experimental Session and Block. The shaded regions in all plots indicate the 83% confidence interval.

## Discussion

The aim of the present work was to assess if 1/*f* aperiodic neural dynamics predict visuomotor performance. Aperiodic resting-state EEG activity is thought to be behaviourally relevant in predicting cognitive capacity (Cross et al., 2020; Ouyang et al., 2020) with higher 1/*f* intercept reflecting higher neural population spiking (Manning et al., 2009; Miller et al., 2012) and steeper 1/*f* slope reflecting a greater excitation-inhibition balance (Lendner et al., 2020; Weber et al., 2020). Despite the apparent association between 1/*f* aperiodic parameters and information processing capacity, to date, there has been no empirical test of these parameters as individual predictors of visuomotor performance. We evaluated 1/*f* intercept and slope as distinct neural markers of information processing capacity and separately evaluated their predictive value for motor and perceptual dimensions of visuomotor performance. Further, we were interested in assessing the predictive value of the aperiodic parameters at increasing levels of visuomotor performance demands as well as across practice with the visuomotor task.

Performance in the motor dimension was indexed with RT, reflecting latencies associated with speeded aiming and trigger pull movements to shoot targets in the virtual reality marksmanship task environment. RT performance was not characterized by individual 1/*f* intercept while individual 1/*f* slope predicted RT performance but only in the intermediate marksmanship task blocks. Specifically, in Blocks 2 and 3 of the marksmanship task, which involved presentation of a target stimulus along with non-target stimuli, RT decreased as individual 1/*f* slope increased. However, 1/*f* slope did not predict RT in the least difficult Block 1 (1 target stimulus only per trial) or RT in Block 4 involving continuous presentation of 2 target stimulus variations and non-target stimuli. The present findings suggest that only the 1/*f* slope predicts individual speeded motor action performance, though the predictive value of 1/*f* slope for RT is dependent on speeded action demands. When motor demands are low, as in Block 1, which had the shortest RT, 1/*f* slope did not predict speeded action. With respect to 1/*f* slope prediction of RT in Block 4, it is not clear if the lack of association is due to high task difficulty or due to changed target stimuli presentation conditions. Although Block 4 was intended to represent highest task difficulty, RT in Block 4 was descriptively shorter than in Block 3. Thus, it is possible that the continuous presentation of the two target stimuli in Block 4 reduced response production demands relative to Block 3. This lowered response costs might explain the lack of RT prediction from 1/*f* slope in the final task block.

Perceptual demands in the marksmanship task were increased between Blocks 2 and 4 based on the number of non-target stimuli presented in a trial as well as through introduction of a novel target stimulus in the final block. Perceptual sensitivity, indexed by d’, across Blocks 2 to 4 confirmed that perceptual demands increased across these task blocks. Both parameters of resting-state aperiodic activity predicted perceptual sensitivity in the marksmanship task. However, the strength of this prediction was most notable in the highest perceptual demand condition. Thus, the predictive value of resting 1/*f* aperiodic parameters for perceptual sensitivity appears to increase as perceptual discrimination demands increase.

Our design included three sessions with the virtual reality marksmanship task, which afforded an opportunity to inspect how prediction of perceptual sensitivity from resting aperiodic neural dynamics is shaped by task experience. The results revealed that practice with the marksmanship task altered 1/*f* intercept and slope associations with perceptual sensitivity performance. In the first session, a higher 1/*f* intercept and a steeper 1/*f* slope were associated with higher d’ in Block 4. This advantage then diminished across the second and third sessions to the extent that individuals with lower a 1/*f* intercept and a shallower 1/*f* slope demonstrated the most gains in Block 4 d’ from practice. Taken together, our findings illustrate three important points about the predictive value of resting aperiodic neural dynamics for perceptual sensitivity. First, 1/*f* intercept and slope appear to distinguish individual perceptual sensitivity capacity only in high perceptual demand conditions. Second, higher 1/*f* intercept and steeper 1/*f* slope predict higher perceptual sensitivity only in initial exposure to high perceptual demand conditions. Finally, a lower 1/*f* intercept and a shallower 1/*f* slope resting-state aperiodic parameters identify individuals who will benefit the most from practice in improving perceptual sensitivity.

The relationship between a steeper 1/*f* slope and enhanced perceptual sensitivity is in keeping with previous work examining the association between aperiodic neural dynamics and cognition (Cross et al., 2020; Ouyang et al., 2020; Ostlund et al., 2021). Here, we demonstrated that both a high 1/*f* intercept (reflective of higher overall neural spiking) and a steeper 1/*f* slope were predictive of enhanced perceptual sensitivity under more difficult conditions of information processing (i.e., Block 4). A steeper 1/*f* slope is posited to represent an increase in neural inhibition activity (i.e., a shift toward more inhibitory states; Gao et al., 2017), and thus increase the consistency of stimulus processing (Tran et al., 2020). By contrast, a shallower slope (i.e., flattening of the aperiodic spectrum) may reflect a diminished neural excitation/inhibition balance and consequently, reduced information processing capacity. For example, children with attention deficit hyperactivity disorder (Ostlund et al., 2021) and older adults (Voytek et al., 2015; Tran et al., 2020) demonstrate shallower 1/*f* slopes and deficits in information processing capacity relative to age-matched healthy controls. Mechanistically, shallower slopes may reflect altered GABAergic and glutamatergic activity in cortical circuitry, neurotransmitters associated with maintaining an optimal excitation/inhibition balance (Ostlund et al., 2021). From this perspective, our finding that a steeper 1/*f* slope is associated with enhanced perceptual sensitivity (d’) lends further support for the argument that a steeper resting aperiodic spectrum slope, and thus a greater inhibitory state, facilitates optimal information processing.

For the 1/*f* intercept, we found that a higher intercept, which is thought to reflect increased neural population spiking (Manning et al., 2009; Miller et al., 2012), predicted enhanced perceptual decision-making, quantified as d’ for the most difficult condition (Block 4) during Session 1. A higher aperiodic intercept has been positively correlated with the fMRI BOLD signal (e.g., Winawer et al., 2013; Jacob et al., 2021). Increases in the BOLD signal are interpreted as increased neural recruitment as oxygenated and deoxygenated hemoglobin concentration changes in response to increased neural metabolism (Glover, 2011). From this perspective, participants with a higher 1/*f* intercept during resting-state periods may have higher basal neural metabolic rates and thereby more available neural capacity (Hall et al., 2016) to facilitate enhanced perceptual decision-making (cf. Ouyang et al., 2020). Regarding initial speeded action, we found that a steeper 1/*f* slope predicted shorter RT in conditions of inter-mediate difficulty (i.e., Blocks 2 – 3) but not under relatively easy or difficult conditions (i.e., Blocks 1 and 4, respectively). This finding suggests that there may be a trade-off in the relationship between the excitation/inhibition balance of cortical circuitry and processing speed under conditions of intermediate difficulty; however, this idea requires further investigation, such as experimentally manipulating (e.g., via the administration of propofol and ketamine; Lendner et al., 2020; Waschke et al., 2021) the excitation/inhibition ratio of relevant neural networks under differing conditions of speeded action demands.

Our specific finding of a relationship between 1/*f* slope (and not intercept) and RT may imply that neural states of higher inhibition, not overall neuronal firing rate, is more relevant to capacity for speeded action. Further, a combined finding relating to both 1/*f* intercept and slope predicting initial perceptual sensitivity, may imply that both overall neural firing rate and the excitation/inhibition balance of neural circuitry are critical for optimal perceptual decision-making. Future work should investigate whether these findings extend beyond resting-state EEG. For example, estimating aperiodic activity during visuomotor learning (e.g., immediately before and during target presentation) may reveal more fine-grained, stimulus-related aperiodic responses related to perceptual decision-making and speeded action (e.g., Cross et al., 2020).

The findings from this initial investigation into the predictive value of resting-state 1/*f* aperiodic neural dynamics for visuomotor performance need to be carefully considered due to some limitations. There is potential that prediction of visuomotor performance with aperiodic parameters of restingstate brain activity is specific to the present virtual reality environment task. For example, the stimuli were presented in a two-dimensional plane and target and non-target stimuli were static. Further, the present marksmanship task included a limited set of conditions intended to challenge both motor and perceptual demands. For example, speeded action was limited to a small range of movement in terms of aiming at targets. In addition, distinction between targets and nontargets was based on differences in helmet shape and blue or red coloring of helmet and armor for cartoon robots. Thus, our demonstration of aperiodic individual differences with a virtual reality visuomotor task might not generalize to performance of natural visuomotor tasks (Kozak et al., 1993; Anglin et al., 2017; Haar et al., 2021). Future work is needed to address if visuomotor performance in more complex naturalistic environments can be predicted by 1/*f* aperiodic neural activity similarly to our reported findings.

There is also the potential that the present findings are limited to healthy young adults. These findings might not generalize to visuomotor performance in younger individuals undergoing both neural (for example see, Miskovic et al., 2015; Anderson & Perone, 2018; Ostlund et al., 2021) and visuomotor (Bo et al., 2006) developmental changes. Moreover, our findings might not extend to older adults given the possibility that aging-related changes in neural (Scally et al., 2018; Tian et al., 2018) and visuomotor (Endrass et al., 2012) function might alter mechanistic relationships between aperiodic neural dynamics and perceptual-motor processes. Thus, future work is needed to investigate the predictive value of resting-state 1/*f* aperiodic neural activity for visuomotor performance across the lifespan. In conclusion, we have demonstrated that individual resting-state 1/*f* aperiodic neural activity predicts motor and perceptual dimensions of visuomotor performance. A steeper 1/*f* slope, thought to represent global levels of neural excitation/inhibition, predicts faster aiming and selection performance and higher perceptual sensitivity in initial exposure to highly demanding visuomotor task conditions. We have also demonstrated that a lower 1/*f* intercept and shallower 1/*f* slope identify individuals who are likely to derive the most gains in perceptual sensitivity from task practice. Aligning the present findings with previous demonstrations of 1/*f* scaling of skilled movement measures highlights the ubiquity of 1/*f* aperiodic dynamics in the neural, cognitive and behavioral dimensions of motor skill performance. These 1/*f* aperiodic dynamics reflect self-organization of coordinative systems that underlie skilled performance (Wijnants et al., 2009).

## ACKNOWLEDGEMENTS

We would like to acknowledge the contribution of Toby Kelty for programming the virtual reality marksmanship task. We also would like to thank Marina Immink for conducting the experiment protocol as research assistant.

The Commonwealth of Australia supported this research through the Australian Army and a Defence Science Partnership agreement with the Defence Science and Technology Group, as part of the Human Performance Research Network. IBS acknowledges the support of an Australian Research Council Future Fellowship (FT160100437).

The authors declare no conflict of interest associated with this work, including its publication.

## Supplementary Material

**Table S1.** Summary of linear mixed-effects model examining the interaction between the Aperiodic Intercept, Session and Block on Reaction Time (milliseconds) to target stimuli.

|  |  |  |  |  |
| --- | --- | --- | --- | --- |
| Formula: Target_RT ~ Aperiodic_Intercept * Session * Block + Age + (1 \| Subject) |  |  |  |  |
| REML criterion at convergence: 5672 |  |  |  |  |
| Scaled residuals |  |  |  |  |
| Min | 1Q | Median | 3Q | Max |
| -2.7393 | -0.5832 | -0.0013 | 0.6107 | 4.5801 |
| Random effects |  |  |  |  |
| Groups | Name | Variance | Std.Dev. |  |
| Subject | (Intercept) | 2387 | 48.86 |  |
| Residual |  | 2634 | 51.32 |  |
| Number of Observations: 536, groups: subjects, 45 |  |  |  |  |
| Fixed effects | Estimate | Std. Error | t value |  |
| (Intercept) | 337.55 | 350.83 | 0.96 |  |
| Aperiodic_Intercept | -17.52 | 15.03 | -1.16 |  |
| session2 | -224.63 | 230.50 | -0.97 |  |
| session3 | 96.35 | 230.38 | 0.41 |  |
| Block.L | -9.07 | 326.68 | -0.02 |  |
| Block.Q | 64.35 | 326.14 | 0.19 |  |
| Block.C | -135.49 | 325.60 | -0.41 |  |
| age | 6.36 | 2.00 | 3.17 |  |
| Aperiodic_Intercept:session2 | -7.48 | 9.87 | -0.75 |  |
| Aperiodic_Intercept:session3 | 7.55 | 9.87 | 0.76 |  |
| Aperiodic_Intercept:Block.L | -13.93 | 14.00 | -0.99 |  |
| Aperiodic_Intercept:Block.Q | 11.68 | 13.97 | 0.83 |  |
| Aperiodic_Intercept:Block.C | -8.37 | 13.95 | -0.60 |  |
| session2:Block.L | 17.56 | 461.49 | 0.03 |  |
| session3:Block.L | 514.65 | 461.14 | 1.11 |  |
| session2:Block.Q | 267.55 | 461.00 | 0.58 |  |
| session3:Block.Q | 43.37 | 460.76 | 0.09 |  |
| session2:Block.C | 479.38 | 460.50 | 1.04 |  |
| session3:Block.C | 422.72 | 460.38 | 0.91 |  |
| Aperiodic_Intercept:session2:Block.L | 2.05 | 19.78 | 0.10 |  |
| Aperiodic_Intercept:session3:Block.L | 23.43 | 19.76 | 1.18 |  |
| Aperiodic_Intercept:session2:Block.Q | 11.07 | 19.75 | 0.56 |  |
| Aperiodic_Intercept:session3:Block.Q | 1.05 | 19.74 | 0.05 |  |
| Aperiodic_Intercept:session2:Block.C | 21.18 | 19.73 | 1.07 |  |
| Aperiodic_Intercept:session3:Block.C | 18.93 | 19.72 | 0.96 |  |

**Table S2.** Summary of linear mixed-effects model examining the interaction between the Aperiodic Slope, Session and Block on Reaction Time (milliseconds) to target stimuli.

|  |  |  |  |  |
| --- | --- | --- | --- | --- |
| Formula: Target_RT ~ Aperiodic_Slope * Session * Block + Age + (1 \| Subject) |  |  |  |  |
| REML criterion at convergence: 5635.2 |  |  |  |  |
| Scaled residuals |  |  |  |  |
| Min<br>-2.7169 | 1Q<br>-0.5755 | Median<br>0.0129 | 3Q<br>0.6113 | Max<br>4.6404 |
| Random effects |  |  |  |  |
| Groups | Name | Variance | Std.Dev. |  |
| Subject | (Intercept) | 2459 | 49.58 |  |
| Residual |  | 2604 | 51.03 |  |
| Number of Observations: 536, groups: subjects, 45 |  |  |  |  |
| Fixed effects | Estimate | Std. Error | t value |  |
| (Intercept) | 766.86 | 147.17 | 5.21 |  |
| Aperiodic_Slope | 8.21 | 61.85 | 0.13 |  |
| session2 | 89.45 | 80.88 | 1.10 |  |
| session3 | -48.40 | 80.57 | -0.60 |  |
| Block.L | 362.32 | 114.78 | 3.15 |  |
| Block.Q | -390.12 | 114.26 | -3.41 |  |
| Block.C | 167.30 | 113.75 | 1.47 |  |
| age | 6.19 | 2.09 | 2.95 |  |
| Aperiodic_Slope:session2 | 67.33 | 39.03 | 1.72 |  |
| Aperiodic_Slope:session3 | 15.23 | 38.87 | 0.39 |  |
| Aperiodic_Slope:Block.L | 22.47 | 55.34 | 0.40 |  |
| Aperiodic_Slope:Block.Q | -87.89 | 55.11 | -1.59 |  |
| Aperiodic_Slope:Block.C | 51.95 | 54.89 | 0.94 |  |
| session2:Block.L | -3.72 | 162.25 | -0.02 |  |
| session3:Block.L | -107.04 | 161.50 | -0.66 |  |
| session2:Block.Q | -91.39 | 161.75 | -0.56 |  |
| session3:Block.Q | 61.06 | 161.14 | 0.37 |  |
| session2:Block.C | -231.72 | 161.26 | -1.43 |  |
| session3:Block.C | -75.53 | 160.77 | -0.47 |  |
| Aperiodic_Slope:session2:Block.L | 12.69 | 78.31 | 0.16 |  |
| Aperiodic_Slope:session3:Block.L | -36.38 | 77.90 | -0.46 |  |
| Aperiodic_Slope:session2:Block.Q | -48.64 | 78.05 | -0.62 |  |
| Aperiodic_Slope:session3:Block.Q | 20.33 | 77.74 | 0.26 |  |
| Aperiodic_Slope:session2:Block.C | -104.85 | 77.80 | -1.34 |  |
| Aperiodic_Slope:session3:Block.C | -27.39 | 77.58 | -0.35 |  |

**Table S3.** Summary of linear mixed-effects model examining the interaction between the Aperiodic Intercept, Session and Block on d’.

|  |  |  |  |  |
| --- | --- | --- | --- | --- |
| Formula: dprime ~ Aperiodic_Intercept * Session * Block + Age + (1 \| Subject) |  |  |  |  |
| REML criterion at convergence: 846.7 |  |  |  |  |
| Scaled residuals |  |  |  |  |
| Min | 1Q | Median | 3Q | Max |
| -3.8414 | -0.4709 | 0.0369 | 0.5917 | 3.3293 |
| Random effects |  |  |  |  |
| Groups | Name | Variance | Std.Dev. |  |
| Subject | (Intercept) | 0.1002 | 0.3165 |  |
| Residual |  | 0.3779 | 0.6148 |  |
| Number of Observations: 403, groups: subjects, 45 |  |  |  |  |
| Fixed effects | Estimate | Std. Error | t value |  |
| (Intercept) | 4.72 | 2.39 | 1.97 |  |
| Aperiodic_Intercept | -0.01 | 0.10 | -0.14 |  |
| session.L | -3.36 | 2.25 | 1.49 |  |
| session.Q | 1.66 | 2.25 | 0.74 |  |
| Block.L | 0.18 | 2.25 | 0.08 |  |
| Block.Q | 1.35 | 2.25 | 0.60 |  |
| age | -0.05 | 0.01 | -3.42 |  |
| Aperiodic_Intercept:session.L | -0.16 | 0.09 | -1.67 |  |
| Aperiodic_Intercept:session.Q | 0.07 | 0.09 | 0.82 |  |
| Aperiodic_Intercept:Block.L | 0.05 | 0.09 | 0.53 |  |
| Aperiodic_Intercept:Block.Q | 0.07 | 0.09 | 0.73 |  |
| session.L:Block.L | -8.82 | 3.89 | -2.26 |  |
| session.Q:Block.L | 6.93 | 3.89 | 1.78 |  |
| session.L:Block.Q | -6.91 | 3.89 | -1.77 |  |
| session.Q:Block.Q | 4.14 | 3.90 | 1.06 |  |
| Aperiodic_Intercept:session.L:Block.L | -0.39 | 0.16 | -2.36 |  |
| Aperiodic_Intercept:session.Q:Block.L | 0.30 | 0.16 | 1.80 |  |
| Aperiodic_Intercept:session.L:Block.Q | -0.29 | 0.16 | -1.77 |  |
| Aperiodic_Intercept:session.Q:Block.Q | 0.18 | 0.16 | 1.08 |  |

**Table S4.**
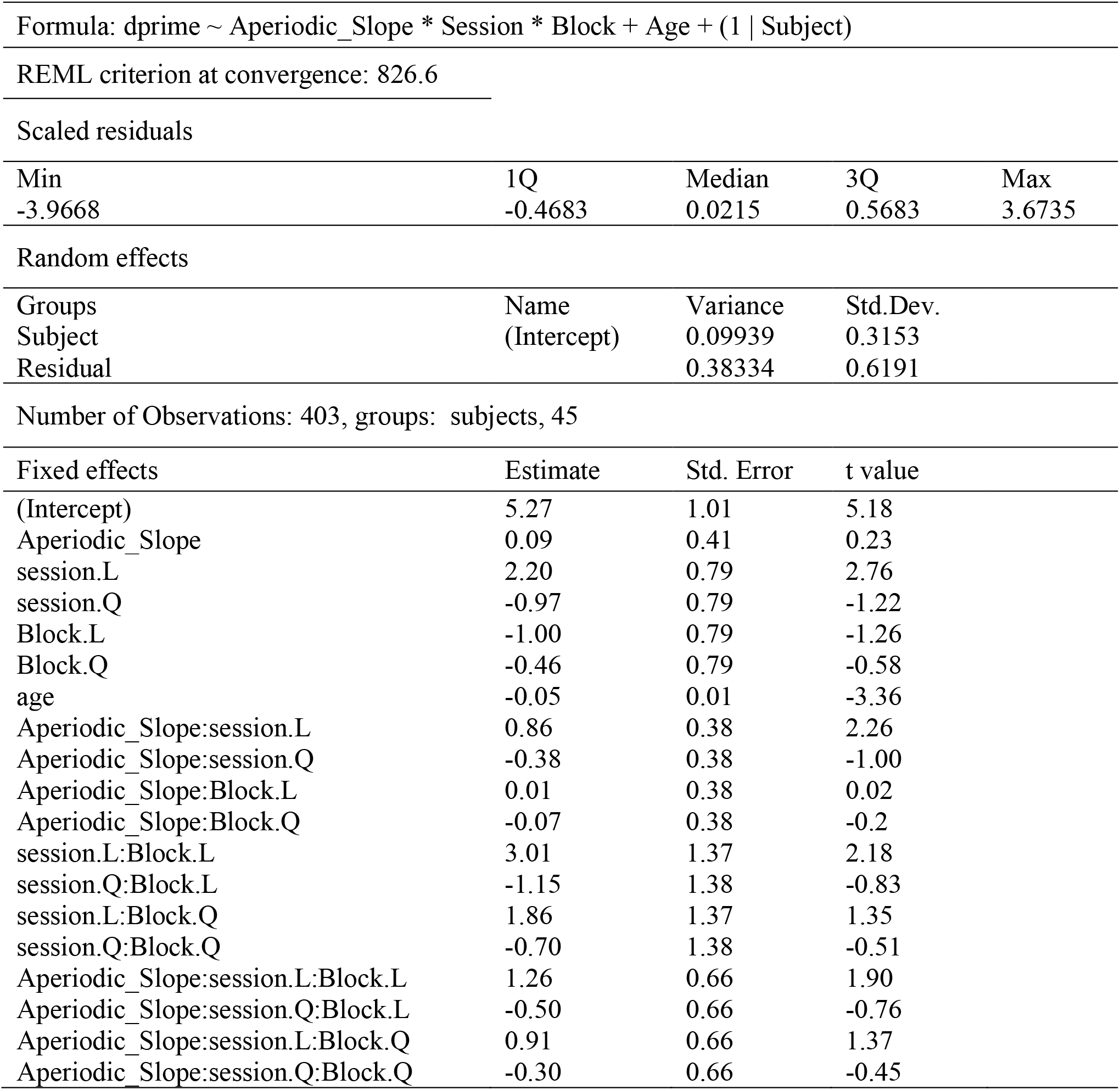
Summary of linear mixed-effects model examining the interaction between the Aperiodic Slope, Session and Block on d’.

|  |  |  |  |  |
| --- | --- | --- | --- | --- |
| Formula: dprime ~ Aperiodic_Slope * Session * Block + Age + (1 \| Subject) |  |  |  |  |
| REML criterion at convergence: 826.6 |  |  |  |  |
| Scaled residuals |  |  |  |  |
| Min | 1Q | Median | 3Q | Max |
| -3.9668 | -0.4683 | 0.0215 | 0.5683 | 3.6735 |
| Random effects |  |  |  |  |
| Groups | Name | Variance | Std.Dev. |  |
| Subject | (Intercept) | 0.09939 | 0.3153 |  |
| Residual |  | 0.38334 | 0.6191 |  |
| Number of Observations: 403, groups: subjects, 45 |  |  |  |  |
| Fixed effects | Estimate | Std. Error | t value |  |
| (Intercept) | 5.27 | 1.01 | 5.18 |  |
| Aperiodic_Slope | 0.09 | 0.41 | 0.23 |  |
| session.L | 2.20 | 0.79 | 2.76 |  |
| session.Q | -0.97 | 0.79 | -1.22 |  |
| Block.L | -1.00 | 0.79 | -1.26 |  |
| Block.Q | -0.46 | 0.79 | -0.58 |  |
| age | -0.05 | 0.01 | -3.36 |  |
| Aperiodic_Slope:session.L | 0.86 | 0.38 | 2.26 |  |
| Aperiodic_Slope:session.Q | -0.38 | 0.38 | -1.00 |  |
| Aperiodic_Slope:Block.L | 0.01 | 0.38 | 0.02 |  |
| Aperiodic_Slope:Block.Q | -0.07 | 0.38 | -0.2 |  |
| session.L:Block.L | 3.01 | 1.37 | 2.18 |  |
| session.Q:Block.L | -1.15 | 1.38 | -0.83 |  |
| session.L:Block.Q | 1.86 | 1.37 | 1.35 |  |
| session.Q:Block.Q | -0.70 | 1.38 | -0.51 |  |
| Aperiodic_Slope:session.L:Block.L | 1.26 | 0.66 | 1.90 |  |
| Aperiodic_Slope:session.Q:Block.L | -0.50 | 0.66 | -0.76 |  |
| Aperiodic_Slope:session.L:Block.Q | 0.91 | 0.66 | 1.37 |  |
| Aperiodic_Slope:session.Q:Block.Q | -0.30 | 0.66 | -0.45 |  |

## Notes

### Competing Interest Statement

The authors have declared no competing interest.

## References

Ackerman, P. L. (1987). Individual differences in skill learning: An integration of psychometric and information processing perspectives. Psychological Bulletin, 102(1), 3–27.

Ackerman, P. L. (1988). Determinants of individual differences during skill acquisition: Cognitive abilities and information processing. Journal of Experimental Psychology: General, 117(3), 288–318.

Ackerman, P. L., & Cianciolo, A. T. (2000). Cognitive, perceptual-speed, and psychomotor determinants of individual differences during skill acquisition. Journal of Experimental Psychology: Applied, 6(4), 259–290.

Adams, J. A. (1957). The Relationship between Certain Measures of Ability and the Acquisition of a Psychomotor Criterion Response. Journal of General Psychology, 56(1), 121–134. doi:10.1080/00221309.1957.9918366

Anderson, A. J., & Perone, S. (2018). Developmental change in the resting state electroencephalogram: Insights into cognition and the brain. Brain and Cognition, 126, 40–52. doi:https://doi.org/10.1016/j.bandc.2018.08.001

Andreska, T., Rauskolb, S., Schukraft, N., Lüningschrör, P., Sasi, M., Signoret-Genest, J., … Sendtner, M. (2020). Induction of BDNF Expression in Layer II/III and Layer V Neurons of the Motor Cortex Is Essential for Motor Learning. The Journal of Neuroscience, 40(33), 6289–6308. doi:10.1523/jneurosci.0288-20.2020

Anglin, J. M., Sugiyama, T., & Liew, S. L. (2017). Visuomotor adaptation in head-mounted virtual reality versus conventional training. Scientific Reports, 7(1), 45469. doi:10.1038/srep45469

Austin, P. C., & Hux, J. E. (2002). A brief note on overlapping confidence intervals. J Vasc Surg, 36(1), 194–195. doi:10.1067/mva.2002.125015

Baetu, I., Burns, N. R., Urry, K., Barbante, G. G., & Pitcher, J. B. (2015). Commonly-occurring polymorphisms in the COMT, DRD1 and DRD2 genes influence different aspects of motor sequence learning in humans. Neurobiology of Learning and Memory, 125, 176–188. doi:https://doi.org/10.1016/j.nlm.2015.09.009

Bates, D., Mächler, M., Bolker, B., & Walker, S. (2015). Fitting Linear Mixed-Effects Models Using lme4. Journal of Statistical Software, 67(1), 1–48. doi:10.18637/jss.v067.i01

Bo, J., Contreras-Vidal, J. L., Kagerer, F. A., & Clark, J. E. (2006). Effects of increased complexity of visuo-motor transformations on children’s arm movements. Human Movement Science, 25(4), 553–567. doi:https://doi.org/10.1016/j.humov.2006.07.003

Cheron, G., Petit, G., Cheron, J., Leroy, A., Cebolla, A., Cevallos, C., … Dan, B. (2016). Brain Oscillations in Sport: Toward EEG Biomarkers of Performance. Frontiers in Psychology, 7(246). doi:10.3389/fpsyg.2016.00246

Christou, A. I., Miall, R. C., McNab, F., & Galea, J. M. (2016). Individual differences in explicit and implicit visuomotor learning and working memory capacity. Scientific Reports, 6(1), 36633. doi:10.1038/srep36633

Clayton, K., & Frey, B. B. (1997). Studies of mental “noise”. Nonlinear Dynamics, Psychology, and Life Sciences, 1(3), 173–180.

Colley, I. D., & Dean, R. T. (2019). Origins of 1/f noise in human music performance from short-range autocorrelations related to rhythmic structures. PLoS One, 14(5), e0216088. doi:10.1371/journal.pone.0216088

Cross, Z. R., Corcoran, A. W., Schlesewsky, M., Kohler, M. J., & Bornkessel-Schlesewsky, I. (2020). Oscillatory and aperiodic neural activity jointly predict grammar learning. bioRxiv. doi:10.1101/2020.03.10.984971

Demuru, M., & Fraschini, M. (2020). EEG fingerprinting: Subject-specific signature based on the aperiodic component of power spectrum. Computers in Biology and Medicine, 120. doi:10.1016/j.compbiomed.2020.103748

Diniz, A., Wijnants, M. L., Torre, K., Barreiros, J., Crato, N., Bosman, A. M. T., … Delignières, D. (2011). Contemporary theories of 1/f noise in motor control. Human Movement Science, 30(5), 889–905. doi:https://doi.org/10.1016/j.humov.2010.07.006

Donoghue, T., Haller, M., Peterson, E. J., Varma, P., Sebastian, P., Gao, R., … Voytek, B. (2020). Parameterizing neural power spectra into periodic and aperiodic components. Nature Neuroscience, 23(12), 1655–1665. doi:10.1038/s41593-020-00744-x

Endrass, T., Schreiber, M., & Kathmann, N. (2012). Speeding up older adults: Age-effects on error processing in speed and accuracy conditions. Biological Psychology, 89(2), 426–432. doi:https://doi.org/10.1016/j.biopsycho.2011.12.005

Fleischman, E. A., & Mumford, M. D. (1989). Abilities as Causes of Individual Differences in Skill Acquisition. Human Performance, 2(3), 201–223. doi:10.1207/s15327043hup0203_4

Fleishman, E. A. (1960). Abilities at Different Stages of Practice in Rotary Pursuit Performance. Journal of Experimental Psychology, 60(3), 162–171. doi:DOI 10.1037/h0044953

Fleishman, E. A. (1972). On the relation between abilities, learning, and human performance. American Psychologist, 27(11), 1017–1032.

Fox, J., & Weisberg, S. (2019). An R Companion to Applied Regression (Third ed.). Thousand Oaks CA: Sage.

Gao, R., Peterson, E. J., & Voytek, B. (2017). Inferring synaptic excitation/inhibition balance from field potentials. Neuroimage, 158, 70–78. doi:10.1016/j.neuroimage.2017.06.078

Gilden, D. L., Thornton, T., Mallon, M. W., & 267. (1995). 1/f noise in human cognition. Science, 267, 1837–1839.

Glover, G. H. (2011). Overview of functional magnetic resonance imaging. Neurosurg Clin N Am, 22(2), 133–139, vii. doi:10.1016/j.nec.2010.11.001

Gramfort, A., Luessi, M., Larson, E., Engemann, D. A., Strohmeier, D., Brodbeck, C., … Hamalainen, M. (2013). MEG and EEG data analysis with MNE-Python. Front Neurosci, 7, 267. doi:10.3389/fnins.2013.00267

Groppe, D. M., Bickel, S., Keller, C. J., Jain, S. K., Hwang, S. T., Harden, C., & Mehta, A. D. (2013). Dominant frequencies of resting human brain activity as measured by the electrocorticogram. Neuroimage, 79, 223–233. doi:https://doi.org/10.1016/j.neuroimage.2013.04.044

Haar, S., Sundar, G., & Faisal, A. A. (2021). Embodied virtual reality for the study of real-world motor learning. PLoS One, 16(1), e0245717. doi:10.1371/journal.pone.0245717

Hall, C. N., Howarth, C., Kurth-Nelson, Z., & Mishra, A. (2016). Interpreting BOLD: towards a dialogue between cognitive and cellular neuroscience. Philos Trans R Soc Lond B Biol Sci, 371(1705). doi:10.1098/rstb.2015.0348

He, B. Y. J. (2014). Scale-free brain activity: past, present, and future. Trends in Cognitive Sciences, 18(9), 480–487. doi:10.1016/j.tics.2014.04.003

He, B. Y. J., Zempel, J. M., Snyder, A. Z., & Raichle, M. E. (2010). The Temporal Structures and Functional Significance of Scale-free Brain Activity. Neuron, 66(3), 353–369. doi:10.1016/j.neuron.2010.04.020

Herszage, J., Dayan, E., Sharon, H., & Censor, N. (2020). Explaining Individual Differences in Motor Behavior by Intrinsic Functional Connectivity and Corticospinal Excitability. Frontiers in Neuroscience, 14(76). doi:10.3389/fnins.2020.00076

Immink, M. A., Verwey, W. B., & Wright, D. L. (2020). The Neural Basis of Cognitive Efficiency in Motor Skill Performance from Early Learning to Automatic Stages. In C. S. Nam (Ed.), Neuroergonomics: Principles and Practice (pp. 221–249). Cham: Springer International Publishing.

Jacob, M., Roach, B., Sargent, K., Mathalon, D., & Ford, J. (2021). Aperiodic measures of neural excitability are associated with anticorrelated hemodynamic networks at rest: a combined EEG-fMRI study. bioRxiv. doi:10.1101/2021.01.30.427861

Judd, C. M., Westfall, J., & Kenny, D. A. (2012). Treating stimuli as a random factor in social psychology: a new and comprehensive solution to a pervasive but largely ignored problem. J Pers Soc Psychol, 103(1), 54–69. doi:10.1037/a0028347

Kozak, J. J., Hancock, P. A., Arthur, E. J., & Chrysler, S. T. (1993). Transfer of training from virtual reality. Ergonomics, 36(7), 777–784. doi:10.1080/00140139308967941

Kyllonen, P. C., & Christal, R. E. (1990). Reasoning ability is (little more than) working-memory capacity? Intelligence, 14, 389–433. doi:10.1016/S0160-2896(05)80012-1

Lendner, J. D., Helfrich, R. F., Mander, B. A., Romundstad, L., Lin, J. J., Walker, M. P., … Knight, R. T. (2020). An electrophysiological marker of arousal level in humans. Elife, 9. doi:10.7554/eLife.55092

Macmillan, N. A., & Creelman, C. D. (1990). Response bias: Characteristics of detection theory, threshold theory, and “nonparametric” indexes. Psychological Bulletin, 107, 401–413.

Manning, J. R., Jacobs, J., Fried, I., & Kahana, M. J. (2009). Broadband Shifts in Local Field Potential Power Spectra Are Correlated with Single-Neuron Spiking in Humans. Journal of Neuroscience, 29(43), 13613–13620. doi:10.1523/Jneurosci.2041-09.2009

Miller, K. J., Hermes, D., Honey, C. J., Hebb, A. O., Ramsey, N. F., Knight, R. T., … Fetz, E. E. (2012). Human Motor Cortical Activity Is Selectively Phase-Entrained on Underlying Rhythms. Plos Computational Biology, 8(9). doi:ARTN e100265510.1371/journal.pcbi.1002655

Mirabella, G. (2014). Should I stay or should I go? Conceptual underpinnings of goal-directed actions. Frontiers in Systems Neuroscience, 8(206). doi:10.3389/fnsys.2014.00206

Miskovic, V., Ma, X., Chou, C.-A., Fan, M., Owens, M., Sayama, H., & Gibb, B. E. (2015). Developmental changes in spontaneous electrocortical activity and network organization from early to late childhood. Neuroimage, 118, 237–247. doi:https://doi.org/10.1016/j.neuroimage.2015.06.013

Ostlund, B. D., Alperin, B. R., Drew, T., & Karalunas, S. L. (2021). Behavioral and cognitive correlates of the aperiodic (1/f-like) exponent of the EEG power spectrum in adolescents with and without ADHD. Developmental Cognitive Neuroscience, 48, 100931. doi:https://doi.org/10.1016/j.dcn.2021.100931

Ouyang, G., Hildebrandt, A., Schmitz, F., & Herrmann, C. S. (2020). Decomposing alpha and 1/f brain activities reveals their differential associations with cognitive processing speed. Neuroimage, 205. doi:ARTN 116304/10.1016/j.neuroimage.2019.116304

Rigoli, L. M., Holman, D., Spivey, M. J., & Kello, C. T. (2014). Spectral convergence in tapping and physiological fluctuations: coupling and independence of 1/f noise in the central and autonomic nervous systems. Frontiers in Human Neuroscience, 8(713). doi:10.3389/fnhum.2014.00713

Scally, B., Burke, M. R., Bunce, D., & Delvenne, J.-F. (2018). Restingstate EEG power and connectivity are associated with alpha peak frequency slowing in healthy aging. Neurobiology of Aging, 71, 149–155. doi:https://doi.org/10.1016/j.neurobiolaging.2018.07.004

Sinanaj, I., Cojan, Y., & Vuilleumier, P. (2015). Inter-individual variability in metacognitive ability for visuomotor performance and underlying brain structures. Consciousness & Cognition, 36, 327–337. doi:10.1016/j.concog.2015.07.012

Swanson, H. L., & McMurran, M. (2018). The impact of working memory training on near and far transfer measures: Is it all about fluid intelligence? Child Neuropsychology, 24(3), 370–395. doi:10.1080/09297049.2017.1280142

Swets, J., Tanner, W. P. J., & Birdsall, T. G. (1961). Decision processes in perception. Psychological Review, 68, 301–340.

Tian, L., Li, Q., Wang, C., & Yu, J. (2018). Changes in dynamic functional connections with aging. Neuroimage, 172, 31–39. doi:https://doi.org/10.1016/j.neuroimage.2018.01.040

Tran, T. T., Rolle, C. E., Gazzaley, A., & Voytek, B. (2020). Linked Sources of Neural Noise Contribute to Age related Cognitive Decline. Journal of Cognitive Neuroscience, 32(9), 1813–1822. doi:10.1162/jocn_a_01584

Vallat, R. (2019). YASA (yet another spindle algorithm): A fast and open-source sleep spindles and slow-waves detection toolbox. Sleep Medicine, 64.

Van Dongen, H. P., Olofsen, E., Dinges, D. F., & Maislin, G. (2004). Mixed-model regression analysis and dealing with interindividual differences. Methods in Enzymology, 384, 139–171. doi:10.1016/s0076-6879(04)84010-2

Van Orden, G. C., Holden, J. G., & Turvey, M. T. (2003). Self-organization of cognitive performance. Journal of Experimental Psychology: General, 132(3), 331–350. doi:10.1037/0096-3445.132.3.331

Veale, J. F. (2014). Edinburgh Handedness Inventory - Short Form: A revised version based on confirmatory factor analysis. Laterality, 19(2), 164–177. doi:10.1080/1357650x.2013.783045

Voytek, B., Kramer, M. A., Case, J., Lepage, K. Q., Tempesta, Z. R., Knight, R. T., & Gazzaley, A. (2015). Age-Related Changes in 1/f Neural Electrophysiological Noise. Journal of Neuroscience, 35(38), 13257–13265. doi:10.1523/JNEUROSCI.2332-14.2015

Waschke, L., Donoghue, T., Fiedler, L., Smith, S., Garrett, D. D., Voytek, B., & Obleser, J. (2021). Modality-specific tracking of attention and sensory statistics in the human electrophysiological spectral exponent. bioRxiv, 2021.2001.2013.426522. doi:10.1101/2021.01.13.426522

Weber, J., Klein, T., & Abeln, V. (2020). Shifts in broadband power and alpha peak frequency observed during long-term isolation. Scientific Reports, 10(1). doi:ARTN 17987/10.1038/s41598-020-75127-0

Welch, M., & Henry, F. M. (1971). Individual Differences in Various Parameters of Motor Learning. Journal of Motor Behavior, 3(1), 78–96.

Wen, H., & Liu, Z. (2015). Separating Fractal and Oscillatory Components in the Power Spectrum of Neurophysiological Signal. Purdue University Research Repository. doi: doi:10.4231/R7WQ01R7

Wen, H., & Liu, Z. (2016). Separating Fractal and Oscillatory Components in the Power Spectrum of Neurophysiological Signal. Brain Topography, 29(1), 13–26. doi:10.1007/s10548-015-0448-0

Wickham, H. (2016). ggplot2: Elegant Graphics for Data Analysis. New York: Springer-Verlag.

Wijnants, M. L., Bosman, A. M., Hasselman, F., Cox, R. F., & Van Orden, G. C. (2009). 1/f scaling in movement time changes with practice in precision aiming. Nonlinear Dynamics, Psychology, and Life Sciences, 13(1), 79–98.

Winawer, J., Kay, K. N., Foster, B. L., Rauschecker, A. M., Parvizi, J., & Wandell, B. A. (2013). Asynchronous Broadband Signals Are the Principal Source of the BOLD Response in Human Visual Cortex. Current Biology, 23(13), 1145–1153. doi:10.1016/j.cub.2013.05.001

Wu, J., Srinivasan, R., Kaur, A., & Cramer, S. C. (2014). Resting-state cortical connectivity predicts motor skill acquisition. Neuroimage, 91, 84–90. doi:10.1016/j.neuroimage.2014.01.026

